# Transient topographical dynamics of the electroencephalogram predict brain connectivity and behavioural responsiveness during drowsiness

**DOI:** 10.1101/231464

**Authors:** Iulia M. Comsa, Tristan A. Bekinschtein, Srivas Chennu

## Abstract

As we fall sleep, our brain traverses a series of gradual changes at physiological, behavioural and cognitive levels, which are not yet fully understood. The loss of responsiveness is a critical event in the transition from wakefulness to sleep. Here we seek to understand the electrophysiological signatures that reflect the loss of capacity to respond to external stimuli during drowsiness using two complementary methods: spectral connectivity and EEG microstates. Furthermore, we integrate these two methods for the first time by investigating the connectivity patterns captured during individual microstate lifetimes. While participants performed an auditory semantic classification task, we allowed them to become drowsy and unresponsive. As they stopped responding to the stimuli, we report the breakdown of frontoparietal alpha networks and the emergence of frontoparietal theta connectivity. Further, we show that the temporal dynamics of all canonical EEG microstates slow down during unresponsiveness. We identify a specific microstate (D) whose occurrence and duration are prominently increased during this period. Employing machine learning, we show that the temporal properties of microstate D, particularly its prolonged duration, predicts the response likelihood to individual stimuli. Finally, we find a novel relationship between microstates and brain networks as we show that microstate D uniquely indexes significantly stronger theta connectivity during unresponsiveness. Our findings demonstrate that the transition to unconsciousness is not linear, but rather consists of an interplay between transient brain networks reflecting different degrees of sleep depth.

**Author summary:** How do we lose responsiveness as we fall asleep? As we become sleepy, our ability to react to external stimuli disappears gradually. Here we sought to understand the rapid fluctuations in brain electrical activity that predict the loss of responsiveness as participants fell asleep while performing a word classification task. We analysed the patterns of connectivity between anterior and posterior brain regions observed during wakefulness in alpha band and showed that this connectivity shifted to slower theta frequencies as participants became unresponsive. We also investigated the dynamics of brain electrical microstates, which represent an alphabet of quasi-stable global brain states with lifetimes of 10-100 milliseconds, and found that the temporal dynamics of microstates slowed down when participants became unresponsive. Using machine learning, we further showed that microstate dynamics prior to a stimulus predict whether subjects will respond to it. We integrated microstates and connectivity for the first time to show that a specific microstate captures connectivity patterns correlated with unresponsiveness during this transition. We conclude that falling asleep is accompanied by a millisecond-level interplay between distinct brain networks, and suggest a renewed focus on fine-grained temporal scales in the study of transitions between levels of consciousness.

## Introduction

As we fall asleep, our brain traverses a series of changes which accompany the loss of sensory awareness and responsiveness to the external world. Despite the subjective ability to classify retrospectively one’s own state as “awake” or “asleep” (Hori et al., 1994), research continues to unravel the gradual transitions happening at behavioural (Ogilvie and Wilkinson, 1984), cellular (Steriade et al., 1993), physiological (Prerau et al., 2014) and cognitive (Goupil and Bekinschtein, 2012) level, starting with early drowsiness and continuing into the deep stages of sleep (Ogilvie, 2001). Characterising these transitions and linking across physiological levels is an important step in the modern attempt to understand access-consciousness (Block, 1996; Koch et al., 2016) and its fluctuations in natural, pathological and pharmacological alterations: sleep (Hobson and Pace-Schott, 2002), disorders of consciousness (Giacino et al., 2014), sedation and anaesthesia (Alkire et al., 2008).

The transition from wakefulness to sleep involves a progressive and sometimes nonlinear loss of responsiveness to external stimuli (Ogilvie and Wilkinson, 1984). Behavioural unresponsiveness does not immediately imply unconsciousness (Overgaard and Overgaard, 2011; Sanders et al., 2013). However, from the perspective of levels of consciousness (Laureys, 2005), the capacity to respond to external stimuli offers an objective measurement in the process of transition between full wakefulness and sleep-induced unconsciousness. The question of how we stop responding to stimuli during drowsiness is related to, but distinct from an investigation of the stages of sleep conventionally defined by specific electrophysiological grapho-elements (Iber et al., 2007; Ogilvie, 2001). Indeed, the loss of responsiveness is and distributed across sleep stages: one study found a rate of unresponsiveness of 28% in stage 1, 76% in stage 2, and 95% in stage 3 of sleep (Ogilvie and Wilkinson, 1984). Here, we are specifically interested in the neural markers that predict our inability to respond as we drift to sleep.

A traditional approach for investigating this question is to look at the changes in EEG spectral power and connectivity, which have been shown to vary across levels of consciousness. During relaxed wakefulness, the EEG of most human subjects is characterised by trains of alpha waves, at around 10 Hz, originating from central-posterior cortical areas (Barry et al., 2007; De Gennaro et al., 2016; Niedermeyer, 2005a). During the early onset of sleep, these alpha oscillations disappear and an alpha rhythm with a different cortical origin (Broughton and Hasan, 1995) emerges in anterior regions (Tanaka et al., 1997), while theta power increases, particularly in central regions (Badia et al., 1994; Niedermeyer, 2005b; Ogilvie, 2001; Wright et al., 1995). Similarly, long-range alpha connectivity disintegrates at the onset of sleep, while lower-frequency theta and delta connectivity increases (Tanaka et al., 2000, 1998; Wright et al., 1995). Several power and connectivity patterns have been associated with the loss of consciousness, sometimes specifically with the loss of responsiveness, such as the anteriorisation of alpha power and connectivity, which has been described in drug-induced loss of responsiveness (Chennu et al., 2016), and frontoparietal connectivity, which has been proposed as a key signature of consciousness (Laureys, 2005) and linked to external awareness (Vanhaudenhuyse et al., 2011). The disruption of frontoparietal connectivity at alpha (8-12 Hz) frequencies has been shown to occur in disorders of consciousness (Chennu et al., 2014) and sedation (Chennu et al., 2016). Although it is still debated whether these are signatures of conscious processing or of processes that almost invariably accompany it (Farooqui and Manly, 2017), brain connectivity patterns currently provide, in practice, useful insights into the transitions between levels of consciousness.

Another method that can be employed to investigate the rapidly changing global state of the brain is that of EEG microstates. A microstate represents a quasi-stable spatial topography of electric field on the scalp (Lehmann, 1990, 1971; Lehmann et al., 1987). The conventional method of analysing microstates in a dataset involves running an unsupervised clustering algorithm on a set of EEG topographies of highest variance, followed by labelling of all EEG samples based on the similarity with the four obtained topographies (Murray et al., 2008; Pasqual-Marqui et al., 1995). Four consistent (Khanna et al., 2014) EEG microstate topographies have been identified in a large population of healthy subjects of all ages during resting-state wakefulness (Koenig et al., 2002) and different microstates have been correlated with different cognitive modalities (Lehmann et al., 2010; Milz et al., 2015; Seitzman et al., 2016), but also with mental disorders, such as narcolepsy (Kuhn et al., 2015). A resting-state study of sleep (Brodbeck et al., 2012) identified four EEG microstate topographies in all stages of sleep nearly identical to those of wakefulness, but occurring with altered temporal parameters. Notably, increased microstate duration was associated with deeper sleep. On the contrary, a different study (Cantero et al., 1999) reported a shorter duration of microstates and suggested a larger repertoire of brain states during the hypnagogic period. Microstates are thought to reflect momentary, global, synchronised (Koenig et al., 2005) networks of the brain, reflecting building blocks of large-scale cognitive processing required for the continuous stream of consciousness (Lehmann, 1990). The neural sources underlying microstates are still being explored (Britz et al., 2010; Milz et al., 2017; Pascual-Marqui et al., 2014). Still, the dynamics of the sequence of microstates itself can be seen as a “syntax” of neural activity that is in and of itself an informative tool for modelling and understanding the rapidly-fluctuating global dynamics of the brain.

Brain connectivity and microstates hence provide complementary perspectives on the neurodynamics underlying the loss of responsiveness as we fall asleep. But what is the relationship between brain networks and microstates? There is evidence that transient brain networks can be resolved in electrophysiological data (Baker et al., 2014; Pascual-Marqui et al., 2014; Vidaurre et al., 2016), but it is an open question whether these networks co-occur with the lifetime of individual microstates. We investigate for the first time how spectral connectivity and EEG microstate dynamics interact as we lose responsiveness during drowsiness. We hypothesise that the spectral changes occurring with the loss of responsiveness mirror those observed in the transition to sleep (Ogilvie, 2001), anaesthesia (Chennu et al., 2016; Purdon et al., 2013) and in disorders of consciousness (Chennu et al., 2014): namely, the disintegration of alpha networks, the loss of posterior alpha power, and the emergence of lower-frequency connectivity and power. Alongside, building on previous research on EEG microstate dynamics during sleep (Brodbeck et al., 2012), we hypothesise similar changes in microstate dynamics accompanying the loss of responsiveness during drowsiness. Finally, given that resting-state network activity is known to fluctuate at millisecond level, we hypothesise that the neural changes in that occur during drowsiness underlie the dynamics of both brain networks and the microstates sequence. Specifically, we investigate the possibility that individual microstates co-occur with distinct transient brain networks, reflecting fleeting changes in the global state of the brain during drowsiness.

To address these questions, we use a subset of data from a previously reported auditory discrimination task where subjects became drowsy and unresponsive (Kouider et al., 2014). The task involved pressing a button corresponding to the classification of the auditory stimulus into one of two categories (object or animal). We obtain five minutes of data before and after the loss of responsiveness due to drowsiness. We first characterise the responsive and unresponsive periods by analysing microstate-blind spectral power and connectivity changes in our dataset. Next, we describe the temporal parameters of EEG microstates during responsiveness and unresponsiveness. To test whether these parameters can reliably predict responsiveness to individual stimuli, we apply machine learning to predict responses and misses to stimuli in our task, based only on pre-stimulus microstate parameters. Finally, we investigate the brain connectivity underlying each of the four canonical microstates after the loss of responsiveness and highlight a previously unknown relationship between spectral connectivity and EEG microstates.

## Methods

### Subjects

Sixteen healthy, native English-speaking, right-handed young adults (mean age = 24, S.D. = 2.75; 6 females) were selected for this experiment out of the eighteen subjects from Experiment 1 in a previous study (Kouider et al., 2014). Two subjects from this dataset were excluded by visual inspection due to a failure to remain asleep for a period longer than five minutes, as assessed using responsiveness to stimuli. The participants were directed to not consume stimulants like coffee and to sleep 1-2 hours less than normally before the experiment. All of the subjects were assessed as easy sleepers on the Epworth Sleepiness Scale (scores 7-14). The participants signed a consent form and were reimbursed for their participation. The experiment was approved by the Cambridge Psychology Research Ethics Committee.

### Experimental procedure

The stimuli consisted of 96 spoken English words chosen from the CELEX lexical database (Linguistic Data Consortium, University of Pennsylvania). Half of the words denoted animals and the other half denoted objects. The subjects were asked to classify each stimulus in its respective category (animal or object) by pressing a button. The stimuli were presented through headphones, with an average distance of 8.4 seconds (minimum 6.2 seconds) between consecutive stimuli, as the subjects were lying with their eyes closed in a reclining chair. To facilitate drowsiness, the task was performed in a dark, acoustically and electrically shielded EEG room, and the participants were told that they could fall asleep at any point during the experiment, although they were asked not to stop responding deliberately while still awake.

### EEG data acquisition

The electroencephalogram was continuously recorded at 500 samples per second from 64 Ag/AgCl electrodes (NeuroScan Labs system) positioned and labelled according to the International 10/20 system, with Cz as a reference and including vertical and horizontal electrooculography channels.

### EEG pre-processing

All analyses that follow were performed using MATLAB scripts (The MathWorks, Inc., Natick, Massachusetts, US). The EEGLAB toolbox (Delorme and Makeig, 2004) was used to facilitate data pre-processing.

The data was filtered between 1 and 40 Hz and the full channel mean was subtracted from each channel for baseline correction. The HEOG and VEOG channels were removed. An Independent Component Analysis (ICA) decomposition was performed using the infomax ICA algorithm (Bell and Sejnowski, 1995). Components capturing ocular or single-channel artefacts were removed from the data by visual inspection and considering the correlation with the HEOG and VEOG channels. An average of 11.6 (S.D. = 8.6) out of 63 components were removed per subject. Channel FT8 was interpolated using spherical interpolation in all subjects due to being noisy in most recordings.

### Data segmentation

We classified responsive and unresponsive periods by inspecting the sequence of hits and misses to individual stimuli. We used a liberal window of 6 seconds to allow for a response to a stimulus, regardless of its correctness. A lack of response within 6 seconds was marked as a miss. The choice of a 6-second window for responsiveness was based on our own pilot studies, where we investigated the longest interval that subjects would make a response during drowsiness in a go task. However, note that most reaction times were below 3 seconds (Fig. 1) and the reaction times increased gradually and later in the task, indicating an increase in drowsiness. This was also established in a previous study on the same data (Kouider et al., 2014).

**Figure 1.**
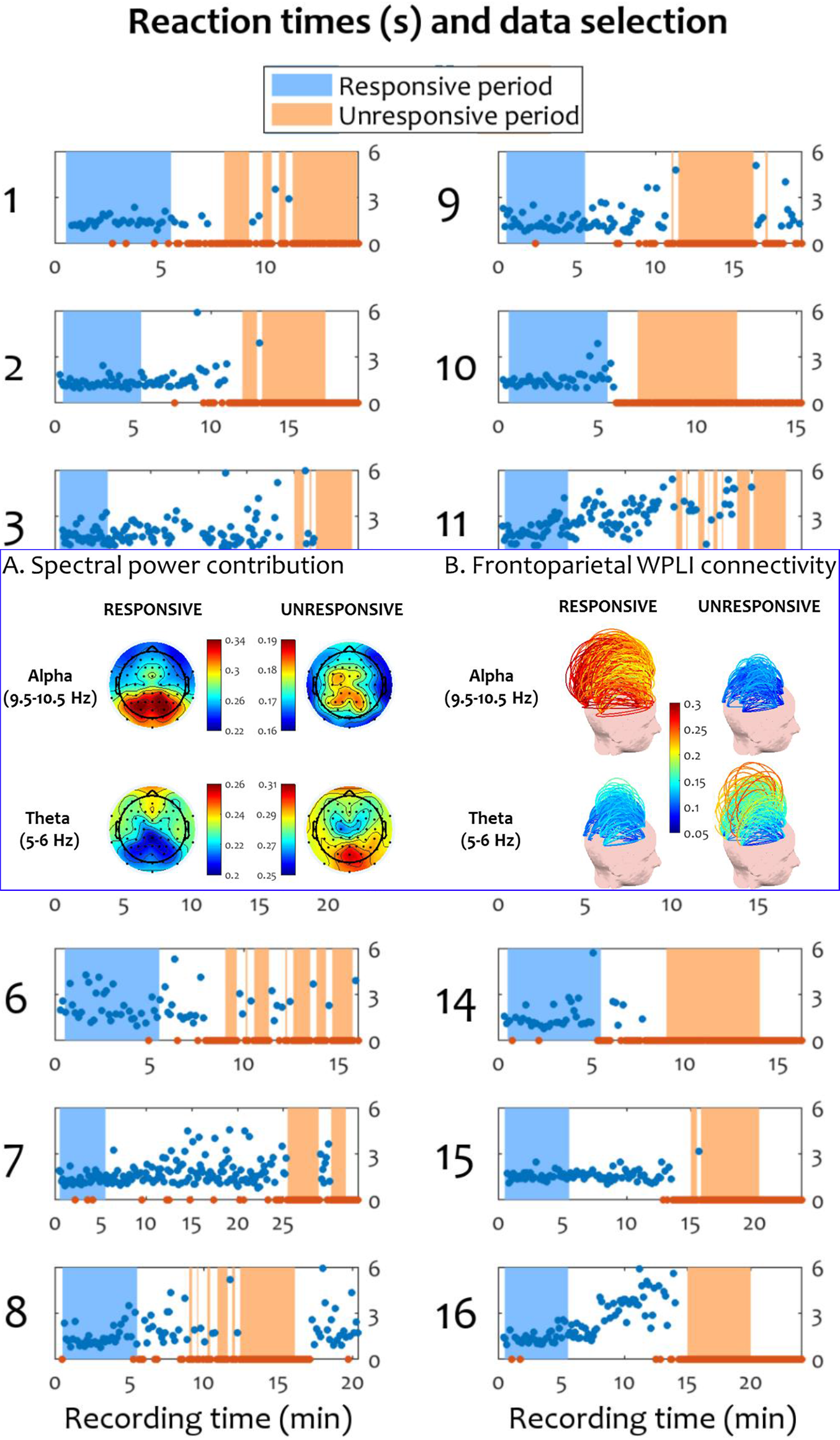
Reaction times and data segmentation into responsiveness and unresponsiveness for individual participants. The horizontal axis represents recording time and the vertical axis represents reaction time in seconds. Blue markers indicate responses, while orange markers indicate misses. The blue area corresponds to the five-minute period of responsiveness, while the orange area corresponds to the five-minute period of unresponsiveness.

For balance across participants and the two behavioural states, a total of five minutes of responsiveness and five minutes of unresponsiveness were extracted from each recording (150000 samples per state, per recording), as shown in Fig. 1. The responsiveness period was taken as the first 0.5 to 5.5 minutes of data in each recording, acquired immediately after the experiment began and the participants were still alert and wakeful. This was confirmed by checking that the large majority of the stimuli were followed by responses during this period; a very small number of misses occurred in more than half of the participants during this period, but they were not contiguous, suggesting task habituation, uncertainty or attention lapses rather than drowsiness-induced unresponsiveness. Then, a period of unresponsiveness was selected by visual inspection of the hits and misses after the end of the responsiveness period, with the aim to find a five-minute interval consisting of as many misses as possible. If a response was present during the period labelled as unresponsiveness, the 10 seconds preceding and following the corresponding stimulus were excluded.

### Microstate topographies

The idea of electric microstates of the brain comes from the observation that the topography of the electric field recorded by EEG over the scalp does not fluctuate randomly, but is instead comprised of short periods of stability (Lehmann, 1971). Four canonical microstates (Koenig et al., 2002), conventionally labelled A, B, C and D, have been shown to be consistent across recording sessions (Khanna et al., 2014) and have been repeatedly confirmed in a wide range of health conditions and cognitive tasks across multiple studies (Britz et al., 2010; Brodbeck et al., 2012; Grieder et al., 2016; Katayama et al., 2007; Kikuchi et al., 2011; Koenig et al., 1999; Kuhn et al., 2015; Milz et al., 2015; Nishida et al., 2013; Pascual-Marqui et al., 2014; Schlegel et al., 2012; Strelets et al., 2003; Tomescu et al., 2014; Van de Ville et al., 2010).

To compute the microstate topographies, the Global Field Power (GFP), representing the standard deviation of the electrode values (Lehmann and Skrandies, 1980), was first computed at each time point. As the number of GFP peaks varied across subjects and condition, we rounded down the minimum number of peaks available and retained the first 5000 peaks in each condition (responsiveness and unresponsiveness) from each recording.

The clustering algorithm was implemented in MATLAB and is presented in Box 1. The algorithm is based on a variant of the method first introduced by (Lehmann et al., 1987), as described in (Murray et al., 2008), and involves an unsupervised clustering of EEG samples into the specified number of classes that best explain the input samples. Note that topographical similarity is computed using the absolute value of the spatial correlation and the polarity of the map is ignored, as topographies with inverted polarities are produced by the same neural generators (Michel et al., 2009). The maximum number of iterations was set to 1000 and the GEV delta was set to 1e-9.

##### Box 1. Microstate clustering algorithm

Microstate clustering algorithm

Input: *n* average-referenced EEG samples (*n* x *number_of_channels*) from GFP peaks.

Output: *k* maps that best characterise the data.

1. Normalize each input sample to a vector of length 1.
2. Pick k random samples as the initial maps.
3. Label each sample as *i* ∈ {1, …*k*}, where *i* is the index of the map with highest absolute spatial correlation.
4. Re-compute each map *i* as the first principal component of each cluster of samples labelled *i*.
5. Compute the Global Explained Variance (GEV).
6. If GEV delta is small enough or maximum number of iterations has been reached, end; else, go to 3.

We initially employed a cross-validation criterion (Pasqual-Marqui et al., 1995) to determine the optimal number of microstates fitting the data, as performed in several previous studies (Brodbeck et al., 2012; Koenig et al., 1999). However, we found that the cross-validation criterion produced different results for when the number of electrodes was down-sampled from 63 to 30 (7 and 4 maps, respectively). This sensitivity of the cross-validation criterion to the number of electrodes has been documented in previous literature (Murray et al., 2008). Hence, we decided to fix the number of microstates to four, in line with previous studies that also fix this number a priori (Khanna et al., 2014; Kikuchi et al., 2007; Koenig et al., 2002; Milz et al., 2015; Schlegel et al., 2012; Strelets et al., 2003; Tomescu et al., 2014).

### Microstate labelling

To obtain the sequence of EEG microstates characterising a recording, each EEG sample was individually assigned to the microstate with the highest corresponding spatial correlation. To correct for noisy assignments during polarity reversals (Koenig and Brandeis, 2016), we applied a previously-described temporal smoothing algorithm for the microstate sequence (Pasqual-Marqui et al., 1995) with parameter b set to 5, corresponding to a smoothing neighbourhood of 20ms. This parameter was chosen to be in the range of mean microstate durations found by (Gärtner et al., 2015) using a model of microstate transition processes based on Markov chains (10 ms during wake, 34 ms during deep sleep).

### Microstate properties

Following the full labelling of each recording, three properties were computed for each microstate per state (responsiveness and unresponsiveness) and per recording:

- The *microstate temporal coverage*, also called the *fractional occupancy*, indicating the percentage of time spend in one microstate;
- The *microstate duration*, indicating the average length of continuous sequences labelled as one microstate;
- The *Global Explained Variance (GEV)*, which measures the amount of spatial correlation of the samples with their corresponding microstate topography, scaled by the GFP.

### Statistics

Interactions between microstate parameters and behavioural state (responsiveness and unresponsiveness) were performed using a two-way repeated measures ANOVA (Hogg and Ledolter, 1987) with the microstate label and the behavioural state as factors. Sphericity was tested using Mauchly’s test of sphericity (Mauchly, 1940) and, where violated, was corrected using the Greenhouse-Geisser procedure (Greenhouse and Geisser, 1959). The Tukey-Kramer method (Tukey, 1949) was used to correct for multiple comparisons. After correction, a conventional threshold of p=0.05 was used to assess significance. Unless otherwise specified, similar statistical tests were also performed for the measures that follow.

### Responsiveness prediction

We applied machine learning classification to explore whether microstate properties identified in the ongoing brain dynamics immediately preceding each auditory stimulus in the experimental trials could predict the presence or absence of a response to that stimulus. All trials were considered for classification, both within and outside the periods labelled as responsive or unresponsive for the above microstate analysis.

Five seconds of EEG data immediately preceding a stimulus were used to generate the features for classification. We also investigated using shorter pre-stimulus time periods, down to 1 second of pre-stimulus data, but we found that classification accuracy increased with a larger amount of prestimulus data over which microstate dynamics could be more accurately estimated. At the same time, the amount of pre-stimulus data was restricted by the overlap with the previous trial. Trials overlapping with a response corresponding to the previous stimulus were excluded. By setting the pre-stimulus window to five seconds, less than 10% of the trials were rejected due to overlap with the previous trial.

The input features generated for classification consisted of either individual microstate parameters computed during the five-second pre-stimulus period in each trial, or a combination of these parameters. The parameters were those we previously characterised at the group level: namely the mean duration, mean coverage, and mean GEV for each microstate separately. The classifier was trained separately with the above individual and combined features. As a baseline, the theta-alpha ratio was also computed for each trial as the ratio between the total power spectral density at 5-6 and 9.5-10.5 Hz respectively, and used as an input feature for the classifier. The classification label for each trial was generated by labelling it as either as a timely response (1) or a miss (0).

We employed leave-one-subject-out cross-validation to test for the generalisability of the classifier’s performance. For this, the data was split into 16 folds, with one fold corresponding to a single participant’s trials. A support vector machine (SVM) (Christianini and Shawe-Taylor, 2000) with a radial basis function kernel (Vert et al., 2004) was trained repeatedly by excluding one fold at the time from the training set and using it as a test set. The SVM was optimised by exhaustive search to use the optimal value for two parameters: the box constraint, which restricts the number of support vectors, and the kernel scale, both in the range [0.001, 1000] in logarithmic steps of 10.

Platt’s method (Platt, 1999) was used to generate class affiliation probabilities from the trained classifier. These continuously varying probabilities were then used to discriminate between responses and misses using both the Receiver Operator Characteristic (ROC) area under the curve (AUC) (Davis and Goadrich, 2006) and the classification accuracy as the percentage of correct predictions out of the total number of predictions. The classification accuracy was also computed by setting the class discrimination threshold as the optimal operating point of the ROC curve and calculating the percentage of correct predictions, using the threshold as a boundary between the two target classes. We used Wilcoxon signed rank tests (Gibbons and Chakraborti, 2011) to probe for significant differences between classification performances.

### Spectral power and connectivity analyses

The spectral power and connectivity during responsiveness and unresponsiveness was investigated in both microstate-blind and microstate-wise analyses. The power spectral density was computed at each EEG sample between 1 and 20 Hz as the absolute value of the Hilbert transform (Marple, 1999) of the bandpass filtered data within windows of 0.25 Hz. We performed most of the analysis on 1 to 20 Hz and focused on theta and alpha power, whose ratio has been shown to track the onset of sleep (Šušmáková and Krakovská, 2007) and has been employed in other studies of drowsiness (Bareham et al., 2014) or impaired consciousness (Lechinger et al., 2013). Within each recording, the spectral power values at each frequency bin were averaged and normalised by the total power within 1 to 20 Hz, thereby obtaining percentages of power contribution at every channel.

The connectivity within each pair of channels was analysed using the Weighted Phase Lag Index (WPLI) (Vinck et al., 2011), a connectivity measure based on the distribution of phase differences between signals designed to correct for volume conduction, which has been previously used to investigate brain connectivity during loss of consciousness (Chennu et al., 2016, 2014; Lee et al., 2013). The WPLI was obtained by pooling over the Hilbert phase of each sample labelled as belonging to a particular microstate.

For both spectral power and connectivity, the median across channels was computed to obtain one value per microstate and frequency of interest.

To further assess topographical changes in connectivity, two sets representing anterior (AFz, Fz, FCz, AF7, AF3, F1, FC1, F3, FC3, F5, F7, AF8, AF4, F2, FC2, F4, FC4, F6, F8) and posterior (CPz, Pz, POz, Oz, P1, P2, PO3, PO4, O1, O2, P3, P5, P7, P4, P6, P8, CP3, CP1, CP2, CP4) electrodes were selected for analysis. Median WPLI connectivity was computed within the anterior and posterior groups separately for each participant.

## Results

### Behavioural data

The distribution of responsiveness and reaction times over time confirmed that all the subjects were responsive for a minimum of six minutes in the beginning of the experimental session and became unresponsive at a later point. During the unresponsiveness period, participants predominantly reached sleep stage N1, and rarely N2, as detailed in (Kouider et al., 2014). Fig. 1 shows the response reaction times and the misses in each participant, in addition to the selection of data for the subsequent microstate analysis. During responsive periods, most subjects had no more than one miss, with a mean of 2.125% of all responses during this period being misses. The grand average of reaction times during the responsive period was 1.5s (S.D. = 0.7).

### Spectral power and connectivity dynamics

Before delving into microstate analyses, we characterised the spectral power and connectivity patterns during responsive and unresponsive periods. We performed a microstate-blind analysis focusing on previously reported changes related to early sleep, but also anaesthesia and disorders of consciousness, including the alteration of posterior, frontal and frontoparietal connectivity. We focused on alpha and theta frequencies, as the theta-alpha ratio has been shown to be the best discriminator between wake and sleep stage 1 (Šušmáková and Krakovská, 2007), however we also confirmed that there were no significant differences in the means of power and median connectivity in beta (12-30 Hz) or gamma (30-40 Hz) between the responsive and unresponsive periods.

Based on the peaks present in alpha and theta bands in our data at 5.5 and 10 Hz (also see Fig. 6 below) and in keeping with canonical definitions of EEG frequency bands, we defined the spectral frequencies of interest in alpha range at 9.5 to 10.5 Hz and the theta frequencies of interest at 5 to 6 Hz, for both power contributions and connectivity.

We observed a decrease in mean alpha power contribution (t(1,15) = 3.34, p = 0.0044, Cohen’s d = 0.83) and an increase in mean theta power contribution (t(1,15) = 7.1, p = 3.5e^−6^, Cohen’s d = 1.77) going from responsiveness and unresponsiveness. As shown in Suppl. Fig. 1, we noted an alpha peak in spectral power present around 10 Hz in the large majority of the participants during the responsive period, which faded during the unresponsive period. Lower-frequency power in the theta frequency range increased during unresponsiveness. A single notable exception was Subject 12, whose alpha peak did not shift into theta range during the unresponsive period, however this subject was preserved in the analysis since there was no evidence that the experiment instructions were not followed. A grand average topographic plot of power at alpha and theta frequencies (Fig. 2A) revealed that the highest alpha power was located in the posterior area during responsiveness. During unresponsiveness, theta power was highest in posterior channels.

**Figure 2.**
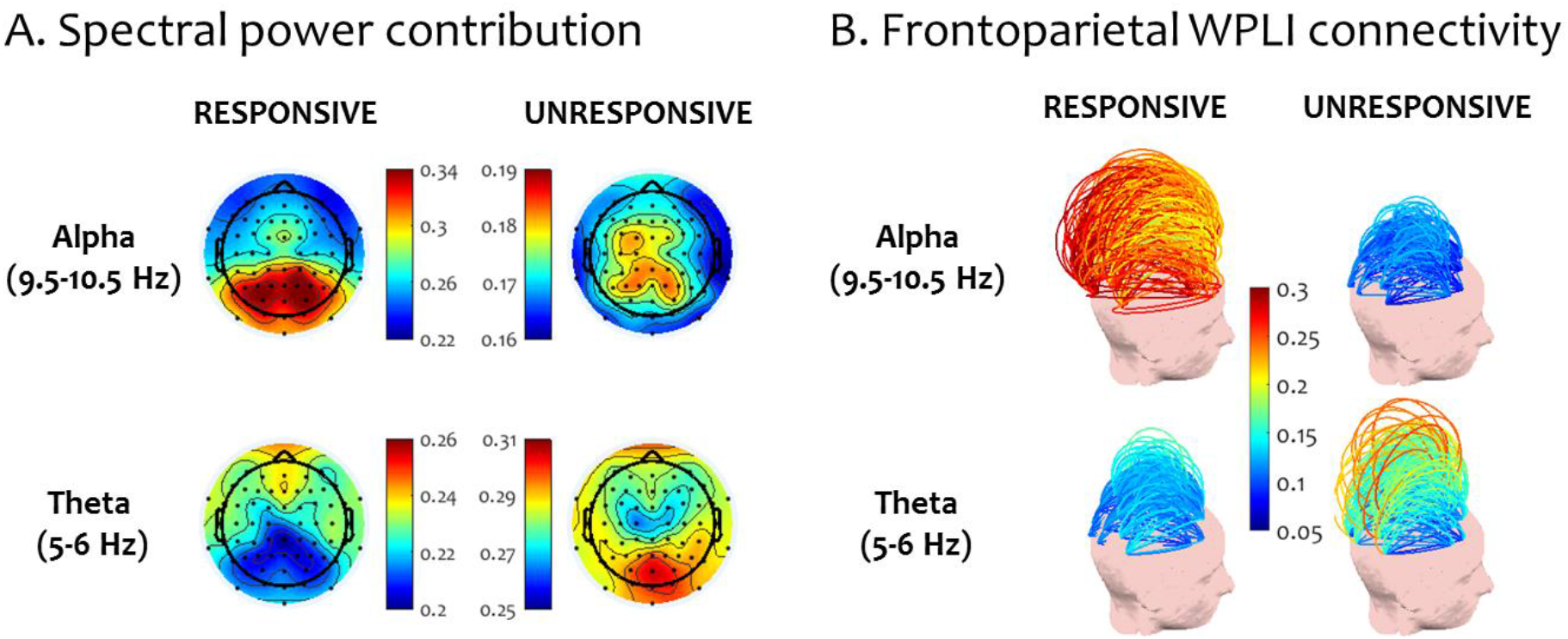
Spectral power topography and WPLI frontoparietal connectivity at alpha (9.5-10.5 Hz) and theta (5-6 Hz) peaks before and after the loss of responsiveness. Values are averaged across participants.

Investigating frontoparietal connectivity in alpha and theta frequencies (Fig. 2B) using the WPLI, we observed the disintegration of long-range connections between frontal and parietal areas going from responsiveness to unresponsiveness at alpha frequencies. A paired t-test confirmed that the median alpha connectivity between the anterior and posterior channels was significantly higher during responsiveness (t(1, 15) = 3.4, p = 0.003, Cohen’s d = 0.85). At the same time, an overall increase in median frontoparietal connectivity was observed in theta frequencies in unresponsiveness, but this was not significant (t(1, 15) = 0.4, p = 0.69, Cohen’s d = 0.1).

### Microstate topographies

It has previously been shown that microstate topographies are highly similar in wakefulness and sleep (Brodbeck et al., 2012). Hence, we applied the microstate clustering algorithm on the set of combined samples from the responsive and unresponsive periods from each subject, in order to obtain four microstate topographies. The resulting maps matched the four canonical microstate topographies commonly described in literature, denoted by letters A to D (Koenig et al., 2002) (Fig. 3). A breakdown of microstate topographies obtained for individual participants is also shown in Suppl. Fig. 3.

**Figure 3.**
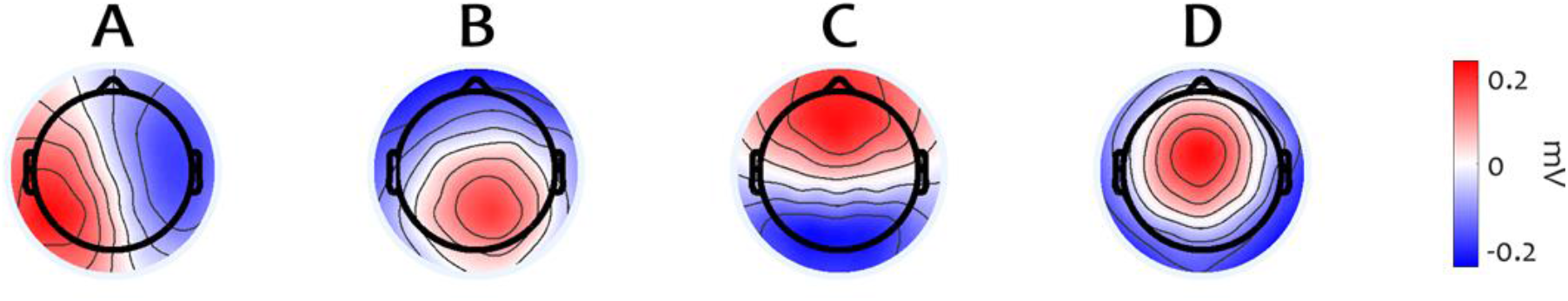
Microstate topographies computed across all subjects.

### Microstate parameters

Having established the topography of the canonical microstates, we next investigated whether the dynamics of the rapid succession of microstates in the EEG remains the same before and after the loss of responsiveness. We computed the duration, the temporal coverage and the global explained variance (GEV) of each microstate during responsiveness and during unresponsiveness (Fig. 4).

**Figure 4.**
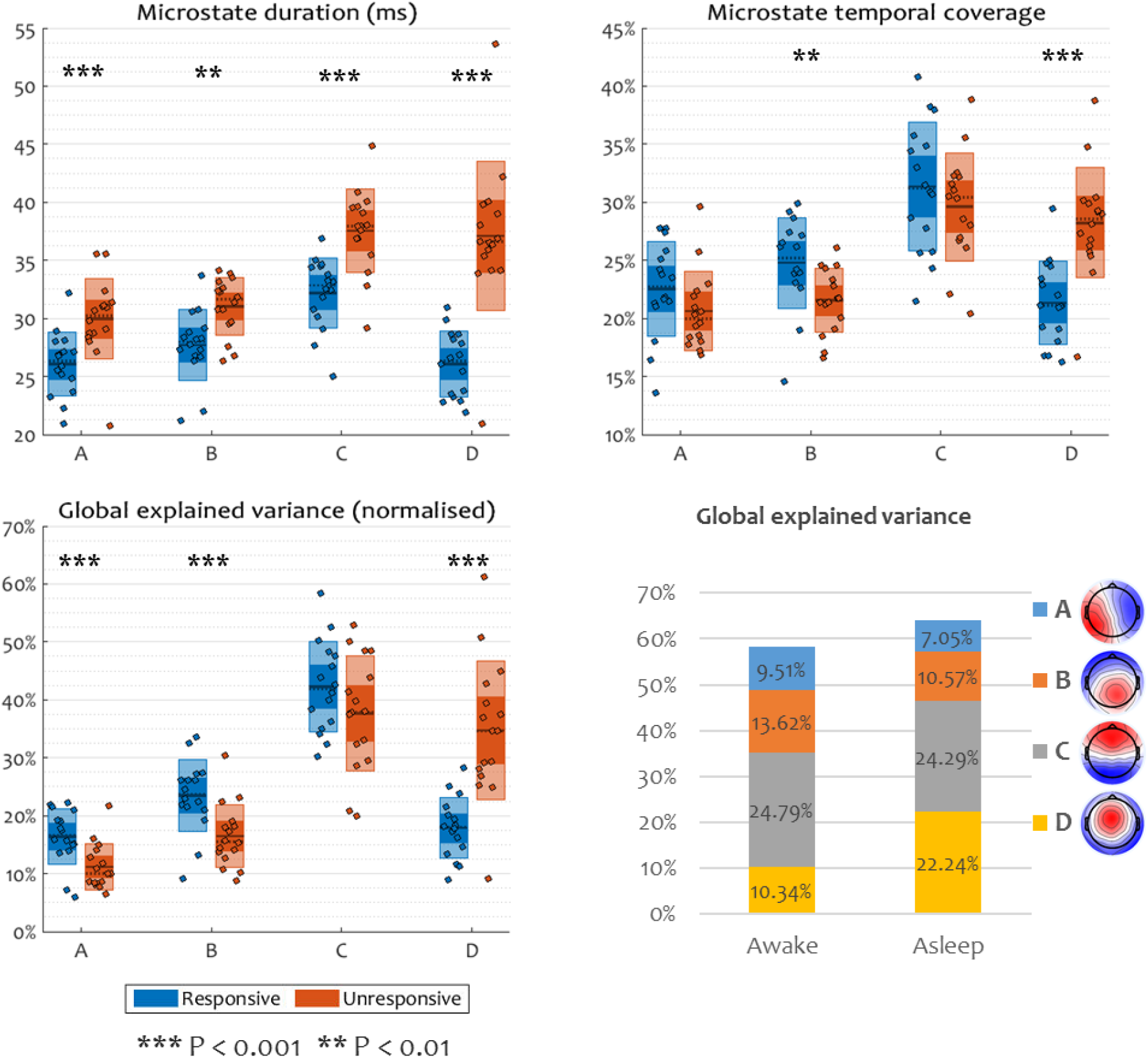
Microstate parameters before and after the loss of responsiveness in drowsiness. Within each group, inner boxes represent the standard error of the mean, outer boxes represent the standard deviation, the mean is shown by a continuous line, the median is shown by a dotted line, and individual participant values are shown as dots. Asterisks show a significant main effect of state within a microstate.

A repeated measures ANOVA with the microstate and the behavioural state (responsiveness and unresponsiveness) as factors found significant interactions between microstate and behavioural state in all of the three microstate parameters investigated: duration (*F_interaction_* = 16.73, *P_interaction_* = 2e^−7^, Cohen’s *d* = 2.11), temporal coverage (*F_interaction_* = 13.08, *P_interaction_* = 3e^−6^, Cohen’s *d* = 1.86) and GEV (*F_interaction_* = 17.95, *P_interaction_* = 8e^−8^, Cohen’s *d* = 2.18). Further exploring the simple effect of state on the parameters within each microstate, the ANOVA revealed that the duration of all microstates was significantly increased during unresponsiveness (*P_state, A_* = 0.0001, *P_state, B_* = 0.003, *P_state, C_* = 0.0001, *P_state, D_* = 3e^−6^), in agreement with previous literature (Brodbeck et al., 2012). Notably, microstate D had a striking increase in duration. At the same time, the temporal coverage of class D was significantly higher during unresponsiveness, whereas the coverage of microstate B was significantly lower during the same period (*P_state, A_* = 0.056, *P_state, B_* = 0.001, *P_state, C_* = 0.26, *P_state, D_* = 1e^−5^). Similarly, the GEV of microstate D was increased during unresponsiveness, while the GEV of microstates A and B were decreased (*P_state, A_* = 0.0002, *P_state, B_* = 0.0002, *P_state, C_* = 0.17, *P_state, D_* = 2e^−5^).

### Single-trial responsiveness prediction

Having characterised the temporal changes in microstate dynamics before and after the loss of responsiveness, we proceeded to verify whether microstate parameters are able to dissociate responsiveness from unresponsiveness at individual trial level during the full recordings, and whether these properties could be generalised across subjects.

Out of all trials, 8% contained a button press event during the five seconds preceding each stimulus and were excluded from further analysis. The remaining data had a balanced distribution of 1078 responses and 1117 misses out of a total of 2195 trials.

Training a radial basis function kernel support-vector machine repeatedly on the combined-microstate and microstate-wise features to predict the binary outcome of a trial, as a response or a miss, using one-subject-out cross-validation, confirmed that microstate dynamics were able to predict responsiveness at individual trial level and across subjects, with a performance similar to that of the established theta-alpha ratio of spectral power (Fig. 5).

**Figure 5.**
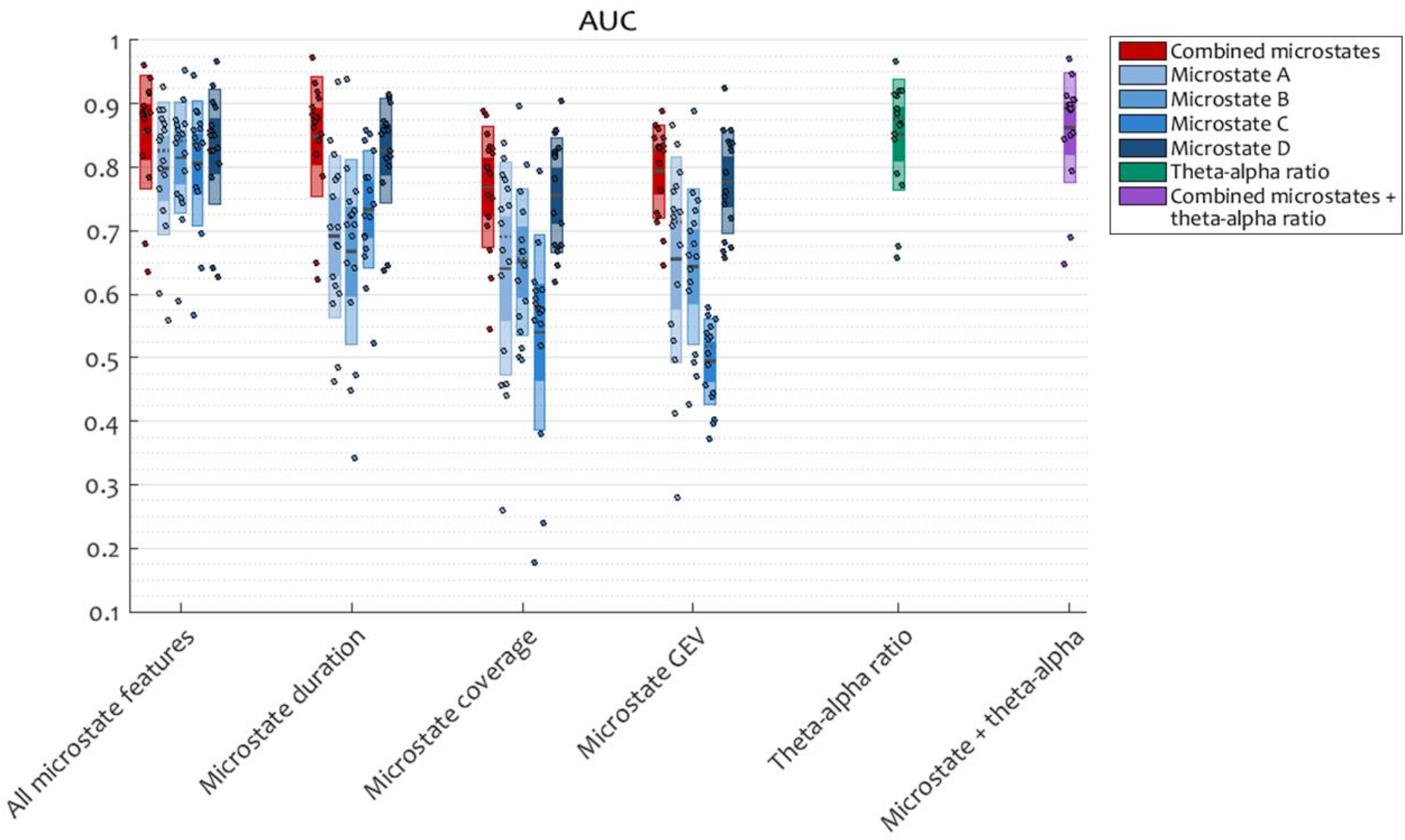
Classification performance, computed as the area under the ROC curve, for a support-vector machine (SVM) trained using 5 seconds of pre-stimulus data to classify responses and misses. Input features are microstate parameters or the theta-alpha ratio, individually or combined. Within each group, inner boxes represent the standard error of the mean, outer boxes represent the standard deviation, the mean is shown by a continuous line, the median is shown by a dotted line, and individual participant values are shown as dots.

Combining the duration, temporal coverage, and GEV of each microstate to obtain a 4 x 5 input feature vector or each trial achieved a mean AUC of 0.8552 (mean classification accuracy of 75.2%). In comparison, the theta-alpha ratio achieved a mean AUC of 0.8519 (mean classification accuracy of 74.24%). A Wilcoxon signed rank test did not find significant differences between these performance distributions. When combined, the microstate features and the theta-alpha ratio obtained a mean AUC 0.8622 (mean classification accuracy of 77.1%).

When used individually as input features for the classification, mean microstate duration performed remarkably well, achieving a mean AUC 0.8484 (mean classification accuracy of 76.1%). According to Wilcoxon test, this was not significantly different from the classification performance of the combined microstate parameters. The duration of microstate D was significantly better at predicting responsiveness than microstates A-C (*p_D-{A,B,C}_*={0.0005, 0.0006, 0.002).

It is worth noting that the one subject for whom the prediction performance was lower in the group was Subject 12, who was also the only one whose alpha peak remained nearly unshifted after the loss of responsiveness (Suppl. Fig. 1).

Taken together, these results indicate that spatiotemporal microstate parameters characterising the pre-stimulus period are indeed informative of the ability of a subject to make a response, similar to the established theta-alpha ratio of the power spectral density. Confirming the initial findings of a more prominent presence of microstate D before the loss of responsiveness due to drowsiness, this microstate also appears to be particularly informative of the capacity of a subject to react to a stimulus. Crucially, these results are generalizable across subjects and valid at single trial level.

### Connectivity differences between microstates

Having established the characteristic temporal patterns exhibited by microstate sequences before and after drowsiness-induced loss of responsiveness, we next proceeded to investigate their relationship with the underlying spectral content of the EEG, and the modulation of this relationship as subjects become unresponsive. To this end, we investigated the power contributions and the WPLI connectivity computed across samples belonging to each microstate before and after the loss of responsiveness. While we do not assume a direct relation between neural sources of EEG microstates and EEG spectral power and connectivity, our aim is to assess whether the neural sources of microstates and sources of spectral measures covary at a fine temporal scale.

The spectral power contribution (Fig. 6A) displayed the characteristic alpha peak around 10 Hz during the responsive period, which faded during the unresponsive period into high power at low frequencies. This pattern was similar during all microstates.

**Figure 6.**
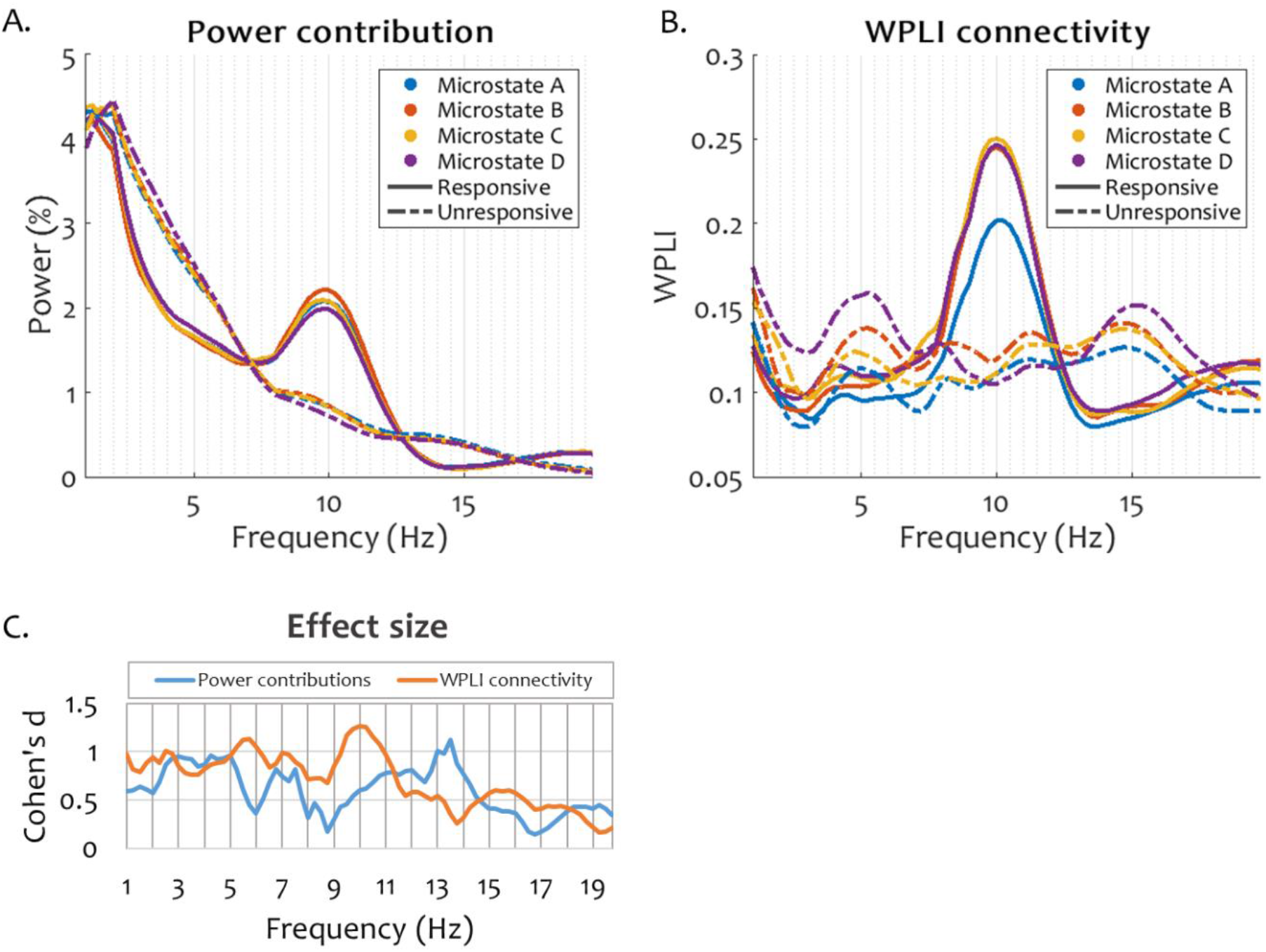
Spectral power contribution (panel A) and WPLI connectivity (panel B) captured during individual microstates before and after loss of responsiveness due to drowsiness. Within each subject, for both power and connectivity, the median across channels was calculated. The figures show the grand average over all subjects. Panel C shows the main effect size, computed as Cohen’s d, of the interaction between behavioural state and microstate at each frequency bin for power contributions and for connectivity.

Likewise, spectral connectivity (Fig. 6B) showed a peak at 10 Hz during responsiveness during all microstates, which faded during unresponsiveness. The only pattern dissociating between microstates during responsiveness was a decreased 10 Hz peak during microstate A. On the other hand, there was a noticeable difference in the level of connectivity during unresponsiveness between all microstate periods, with microstates D and A exhibiting the highest and the lowest connectivity, respectively.

The effect size of the interaction between microstate and behavioural state (responsiveness and unresponsiveness) computed individually at each frequency was indeed generally higher in connectivity than in power (Fig. 6C). The effect size was largest in connectivity at 5.5 Hz and 10 Hz, corresponding to the theta and alpha peaks displayed during all microstates during the unresponsive and responsive periods, respectively. A peak in power contribution was also found at 13.5 Hz, potentially due to the emergence of sleep spindles at the onset of sleep.

We also attempted to use pre-stimulus WPLI connectivity levels at alpha and theta frequencies in order to train a classifier to predict responsiveness, using the same procedure as for the microstate spatiotemporal parameters. Intriguingly, no classifiers could be obtained that exceeded a 60% mean accuracy, either microstate-wise or on the full set of pre-stimulus samples.

### Connectivity during microstate D after the loss of responsiveness

Gathering from the evidence of increased temporal presence of microstate D after the loss of responsiveness, as well as the higher connectivity displayed during this microstate during unresponsiveness in comparison with the microstates A-C, we next sought to understand the spectral connectivity patterns captured during microstate D in the selected alpha and theta ranges during the unresponsiveness period.

Preliminary assessments of connectivity patterns during the four microstates during unresponsiveness revealed visual differences in anterior and posterior connectivity during microstate D as compared to microstates A-C. Considering previous literature (Morikawa et al., 1997; Tanaka et al., 2000, 1998; Wright et al., 1995) suggesting that key changes in connectivity related to the onset of sleep occur topographically in anterior and posterior scalp regions of interest (ROI), as well as frontoparietal having been proposed as a key signature of consciousness (Laureys, 2005), we decided to investigate the within-anterior, within-posterior and between anterior-posterior connectivity during microstate D in comparison with microstates A-C. For this purpose, we performed three repeated measures ANOVA tests to compare the median connectivity during microstate D and that during each of the microstates A-C in each of the six conditions (two frequency bands X three scalp ROIs) during the unresponsive period. Within each condition, we corrected for the false discovery rate across the three tests (D vs A, D vs B and D vs C) using Storey’s procedure (Storey, 2002).

Fig. 7 exemplifies the most prominent differences we found in connectivity between samples covered by microstate D and microstates A-C respectively, during unresponsiveness.

**Figure 7.**
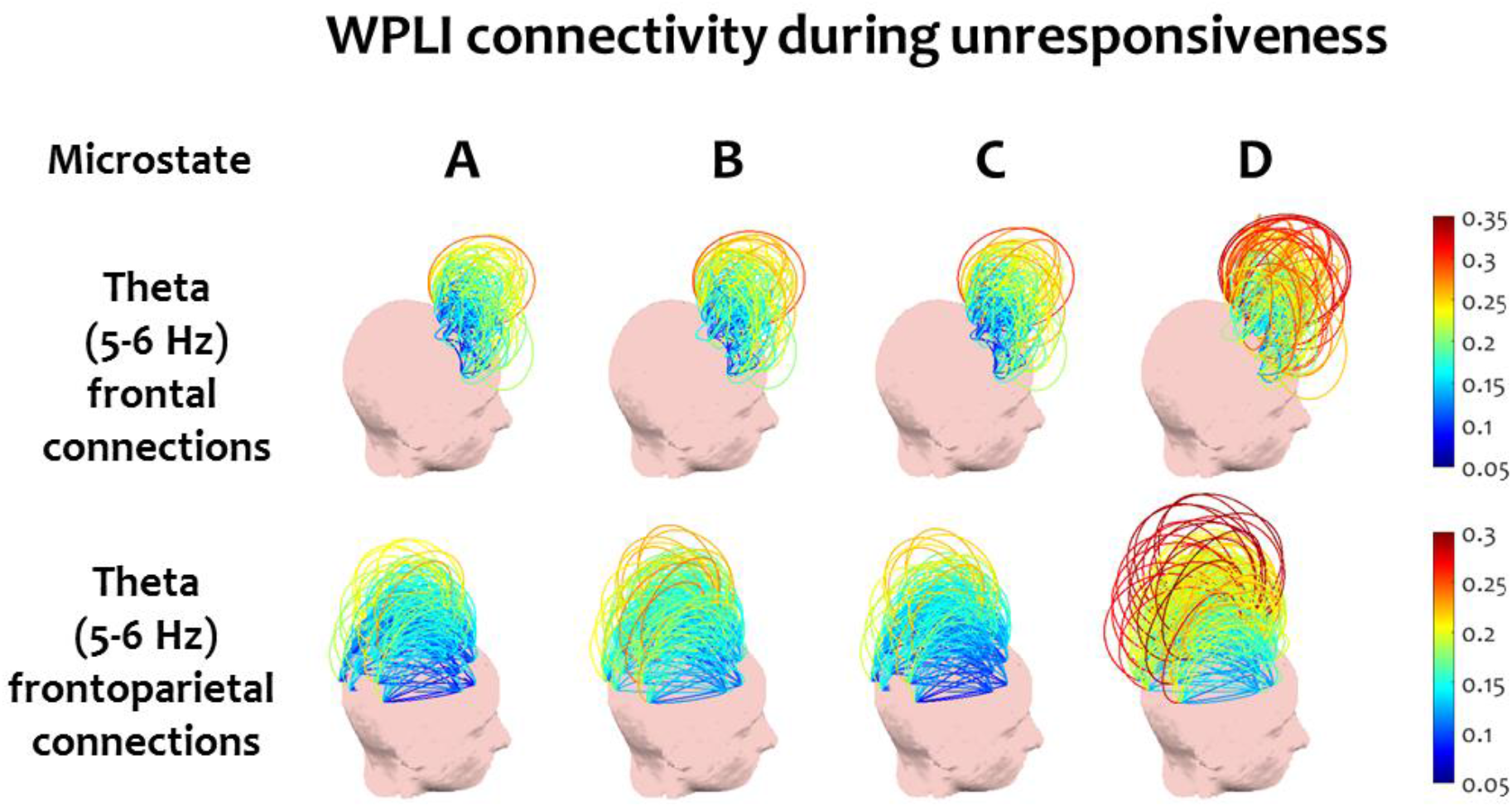
Frontal and frontoparietal WPLI connectivity at theta peak (5-6 Hz). Microstate D captures significantly higher connectivity in these examples compared to microstates A-C.

At the selected theta peak, the t-test results showed significantly higher median connectivity within the anterior region during microstate D compared to each of the other microstates (P_D-{A,B,C}_ = {0.001, 0.008, 0.001}, t_D-{A,B,c}_ = {3.958, 3.069, 4.088}, Cohen’s d_D-{A,B,c}_={0.990, 0.767, 1.022}). Median connectivity between the anterior and posterior regions was also significantly higher during microstate D than in microstates A and C (P_D-{A,B,C}_ = {0.003, 0.297, 0.003}, t_D-{A,B,C}_ = {3.578, 1.081, 3.392}, Cohen’s d_D-{A,B,c}_={0.894, 0.27, 0.848}). No significant differences were found in median connectivity within the posterior area.

Conversely, at the selected alpha peak, the repeated measures ANOVA showed significantly lower median connectivity within the posterior area during microstate D compared to microstates A-C (P_D-{A,B,C}_ = {0.033, 0.037, 0.033}, t_D-{A,B,c}_ = {2.686, 2.294, 2.559}, Cohen’s d_D-{A,B,c}_={0.672, 0.573, 0.67}). At the same time, microstate D captured significantly higher within-anterior median connectivity than microstate A (P_D-{A,B,C}_ = {0.043, 0.617, 0.055}, t_D-{A,B,c}_ = {2.769, 0.511, 2.297}, Cohen’s d_D-{A,B,c}_={0.692, 0.128, 0.574}). No significant difference in median connectivity between anterior and posterior regions was found during microstate D compared to microstates A-C.

These results confirmed that the timecourse of microstate D uniquely capture a simultaneous disintegration of posterior alpha connectivity and emergence of frontal theta connectivity, which is associated with the suppression of responsiveness at the onset of sleep.

## Discussion

### Summary

In this study, we used high-density EEG to explore the transient spatiotemporal and spectral dynamics of electrical brain activity before and after the loss of behavioural responsiveness due to drowsiness. Importantly, we examined the loss of responsiveness as we fall asleep, as opposed to an investigation of canonical sleep stages. Here, unresponsiveness – the failure to respond to the auditory cues elicited by increased drowsiness – provided an objective and non-invasive behavioural criterion in the transitional stage in between full wakefulness and early sleep.

We began by showing differences in spectral power and connectivity after the loss of responsiveness that have been previously shown to differentiate between healthy wakefulness and sleep, sedation and disorders of consciousness: a decrease in posterior alpha power and the emergence of theta power, as well as the disintegration of frontoparietal connectivity in alpha band. We then characterised the spatiotemporal parameters of the four canonical EEG microstates before and after the loss of responsiveness. We showed that microstate parameters not only correlate with behaviour at the group level, but also predict behaviour at the level of individual experimental trials. The ongoing microstate dynamics, particularly the properties of microstate D, before the onset of an auditory stimulus in an experimental trial significantly predicted the likelihood of a response to that auditory stimulus as participants transitioned towards sleep. Specifically, when microstate D occurred more often during the pre-stimulus period, participants were less likely to generate a response to the subsequent stimulus. This relationship highlights a possible functional role of this microstate in modulating behaviour, and the predictive power of this signature to define the capacity to consciously respond to abstract/semantic stimuli. Finally, we examined the spectral power and connectivity characteristics captured during the lifetimes of the four canonical EEG microstates. We discovered that while the distribution of spectral power remains the same across the temporal microstates, spectral connectivity has distinct profiles. We showed that this non-uniform pattern of connectivity across microstates is modulated specifically after the loss of responsiveness: the timecourse of microstate D captured significantly increased connectivity in the theta band after the loss of responsiveness, underpinning a novel profile of interaction between the temporal sequence of microstates and spectral brain connectivity.

### Alpha power and connectivity characterise responsive wakefulness

Our analysis of EEG connectivity before microstate segmentation strengthens the evidence for the fundamental role of the frontoparietal alpha networks in sustaining a state of responsive wakefulness (Laureys, 2005). Alpha band frontoparietal connections have also been shown to disintegrate in disorders of consciousness (Chennu et al., 2014) and sedation (Chennu et al., 2016). Importantly, it is not the full disappearance of all frontoparietal connectivity that drives the loss of responsiveness, but specifically connectivity at alpha frequency. Indeed, literature shows that connectivity shifts from alpha into lower-frequency theta and delta frequencies as consciousness fades (Chennu et al., 2016, 2014; Ogilvie, 2001; Tanaka et al., 2000, 1998; Wright et al., 1995). In the larger picture of states and levels of consciousness, our findings confirm long-range alpha networks as a common marker of consciousness, whether this impairment is natural (sleep), pathological (disorders of consciousness) or pharmacological (sedation).

### Microstate D predicts responsiveness across subjects

Upon examining the spatiotemporal parameters of the canonical EEG microstates, we found an increase in temporal coverage after the loss of responsiveness uniquely specific to microstate D, along with an increase in its global explained variance, as compared to responsive periods. While the duration of all microstates was longer during unresponsiveness, the duration of microstate D had a prominent relative increase. In contrast, the temporal coverage of microstate B decreased in the unresponsive period, as did the global explained variance of microstates A and B. Further, we demonstrated that pre-stimulus parameters of EEG microstate sequences are indeed informative of the capacity of a subject to respond to a stimulus during drowsiness at individual trial level. Again, the special significance of microstate D during unresponsiveness was visible from its increased ability to predict the likelihood of a response, in comparison with microstates A-C. In addition, we showed that the increase in duration of this microstate is the best predictor of responsiveness among all the microstate parameters.

Our usage of machine learning allows us to quantify the performance of the model using its discrimination accuracy, which speaks for the real-world applicability of the method (Breiman, 2001). Moreover, one-subject-out cross-validation allows us to infer that these results are generalizable across people. At the same time, as expected, individual variability caps the maximum possible accuracy when predicting responsiveness. Our results suggest that this cap is around an accuracy of 75% (mean AUC around 0.85). Interestingly, the theta-alpha ratio, which we used as a baseline given its sensitivity as a sleep index (Šušmáková and Krakovská, 2007), achieved a similar classification accuracy as the microstate-based input features. Intriguingly, we were not able to use frontoparietal connectivity as a feature to train a suitable classifier for responsiveness during drowsiness, either considering or ignoring the microstate sequence, despite strong evidence of major connectivity changes occurring before and after the loss of responsiveness. This suggests that connectivity better predicts the level of consciousness estimated over longer time scales, whereas spatiotemporal microstate dynamics capture short-term changes in brain state that predict responsiveness.

### Microstate D captures a distinct connectivity profile after loss of responsiveness

Alongside the distinctive increase in temporal coverage and duration of microstate D, we found a singular spectral connectivity pattern during this microstate after loss of responsiveness, indicating increased median connectivity in theta band, particularly in frontal and frontoparietal connections. At the same time, median posterior connectivity during microstate D was reduced during unresponsiveness. Hence, the timecourse of microstate D appears to uniquely capture a connectivity pattern specific to deeper stages of sleep, in comparison with other microstates present during the same sleep stage. (Britz et al., 2010) have previously reported the lack of any interaction between temporal microstates of the brain and the spectral power of its oscillations, i.e, the spectral power profiles of EEG microstates do not differ from each other, a finding which we replicated. In contrast, we have shown that spectral connectivity presents a significant interaction with temporal microstates, underpinned by the connectivity captured by microstate D.

There currently exists no consensus on the meaning of individual microstates in terms of their neural generators. However, microstate D has occasionally been linked to attentional networks. In a study of BOLD resting-state networks, (Britz et al., 2010) showed a higher correlation of microstate D with ventral and dorsal frontal-parietal networks, functionally associated with attention switching and directing attention towards external salient stimuli. A decreased duration of this microstate has been reported in schizophrenia (Koenig et al., 1999; Lehmann et al., 2005; Nishida et al., 2013; Tomescu et al., 2014) and hallucination (Kindler et al., 2011) - two conditions involving impairments in task switching and attention (Collerton et al., 2005; Cornblatt and Keilp, 1994). An investigation of modalities of thinking found an increased microstate D duration in resting-state compared to visual and verbal task periods (Milz et al., 2015); this was also interpreted as a confirmation of the previously-mentioned study by (Britz et al., 2010) due to a higher probability of attention switching during rest (high microstate D duration), as opposed to performing a single goal-oriented task (lower microstate D duration). On the other hand, (Seitzman et al., 2016) have found an increased duration of microstate D during a cognitive task as compared to wakeful rest.

Given the weak evidence in the literature associating microstate D with task-related attention networks, we are cautious in interpreting our findings on this basis. A previous study on the same data (Kouider et al., 2014) found that a correct response to stimuli is still prepared during unresponsiveness, suggesting preserved attention. It is possible that our findings indicate more demand from attention networks as drowsiness increases and subjects become unable to respond to the task. In study of microstates during sleep in the absence of any task, (Brodbeck et al., 2012) did not observe an increase in this microstate during sleep. This suggests that microstate D might indeed be specifically related to the experimental task. Further, this interpretation is compatible with a study by Katayama et al. (Katayama et al., 2007), which found that the duration of microstate D was increased in light (but not deep) hypnosis, a state which produces similar EEG patterns to sleep-induced unresponsiveness (Barker and Burgwin, 1949).

Nonetheless, the spatiotemporal and spectral connectivity dynamics observed in microstate D after the loss of responsiveness yield an important insight into the dynamics of the transition to sleep. While connectivity averaged during all microstates reflects typical changes commonly found in the loss of consciousness in the onset of sleep, anaesthesia or disorders of consciousness – weaker alpha and stronger theta long-range networks – the individual timecourse of microstate D captures significantly stronger patterns, despite having a duration no longer than 40ms. This suggests that, after the loss of responsiveness, the process of falling asleep is not necessarily linear, but rather consists of an interplay between distinct networks, captured by different microstates, which at are different points along the transition between wakeful and asleep modes of operation. This finding might lend itself to explaining one of the current riddles of sleep research: why is it that, despite the establishment of a series of clear EEG markers delimiting wake and several stages of sleep, finding an EEG-based threshold to separate between the subjective intuition of being awake or asleep has not yet been achieved? Indeed, it has been reported by Hori et al. (1994) that 26% of all subjects assess that they had been awake at times when their EEG was classified as stage 2 sleep, which is commonly used to define “true sleep” (Ogilvie, 2001). The rapid fluctuation of brain networks, some of which are closer to wakefulness (during microstates A-C) and others closer to sleep (during microstate D) could be the reason why our momentary introspective state of being “awake” and “asleep” might not concur with a coarse-grained assessment of EEG over many seconds of data, as usually done during the identification of sleep stages. Instead, our findings here highlightx that further research should focus on the rapidly changing dynamics of brain networks that appear to capture key dynamics relevant to our behavioural and perhaps even introspective state, as we drift into unconsciousness.

## Acknowledgements

We thank Louise Goupil for collecting the data for this experiment.

## Supporting figures

**Supplementary Figure 1.**
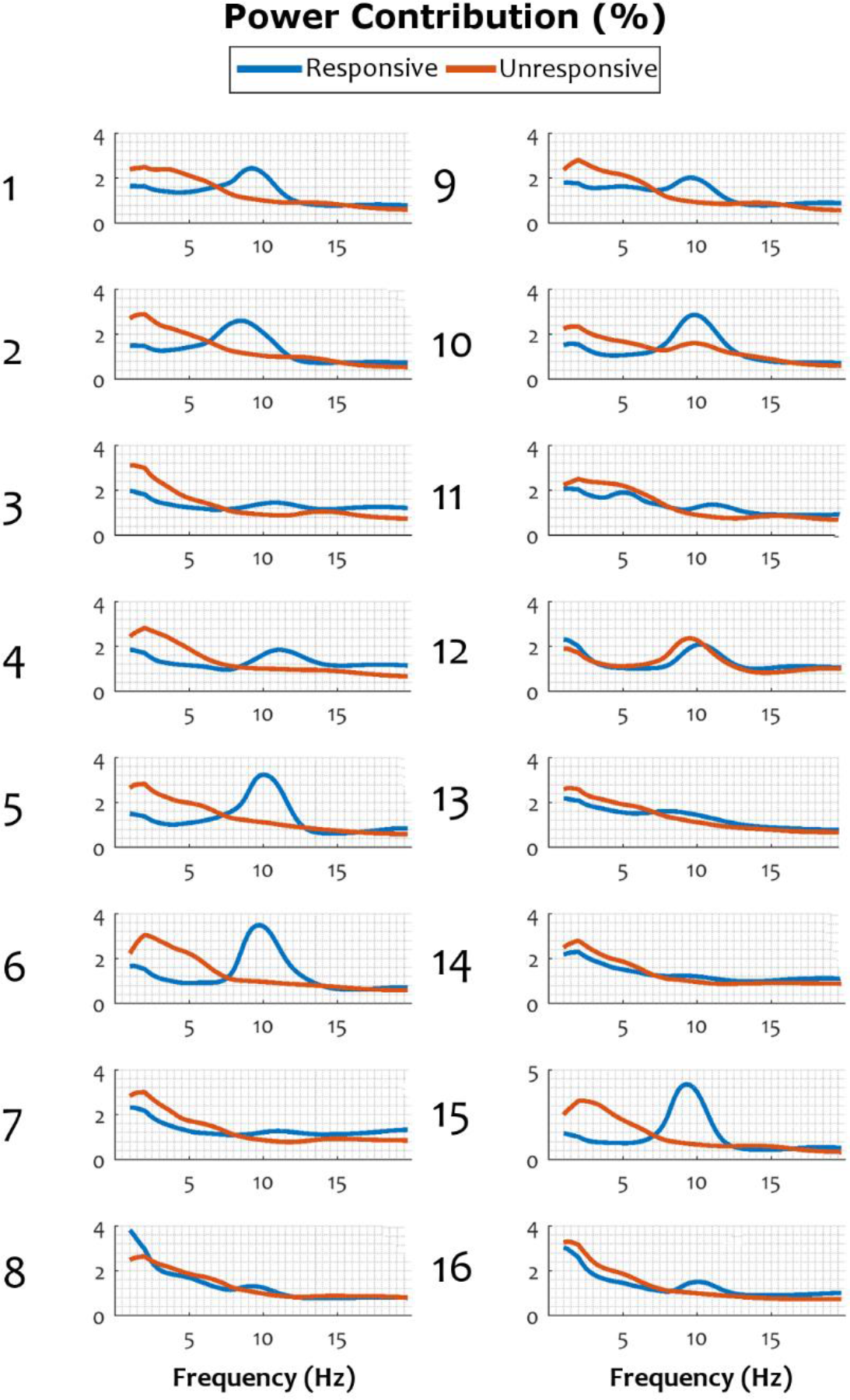
Individual subject spectral power contributions before and after loss of responsiveness. For each subject, values are averaged over posterior channels (see text).

**Supplementary Figure 2.**
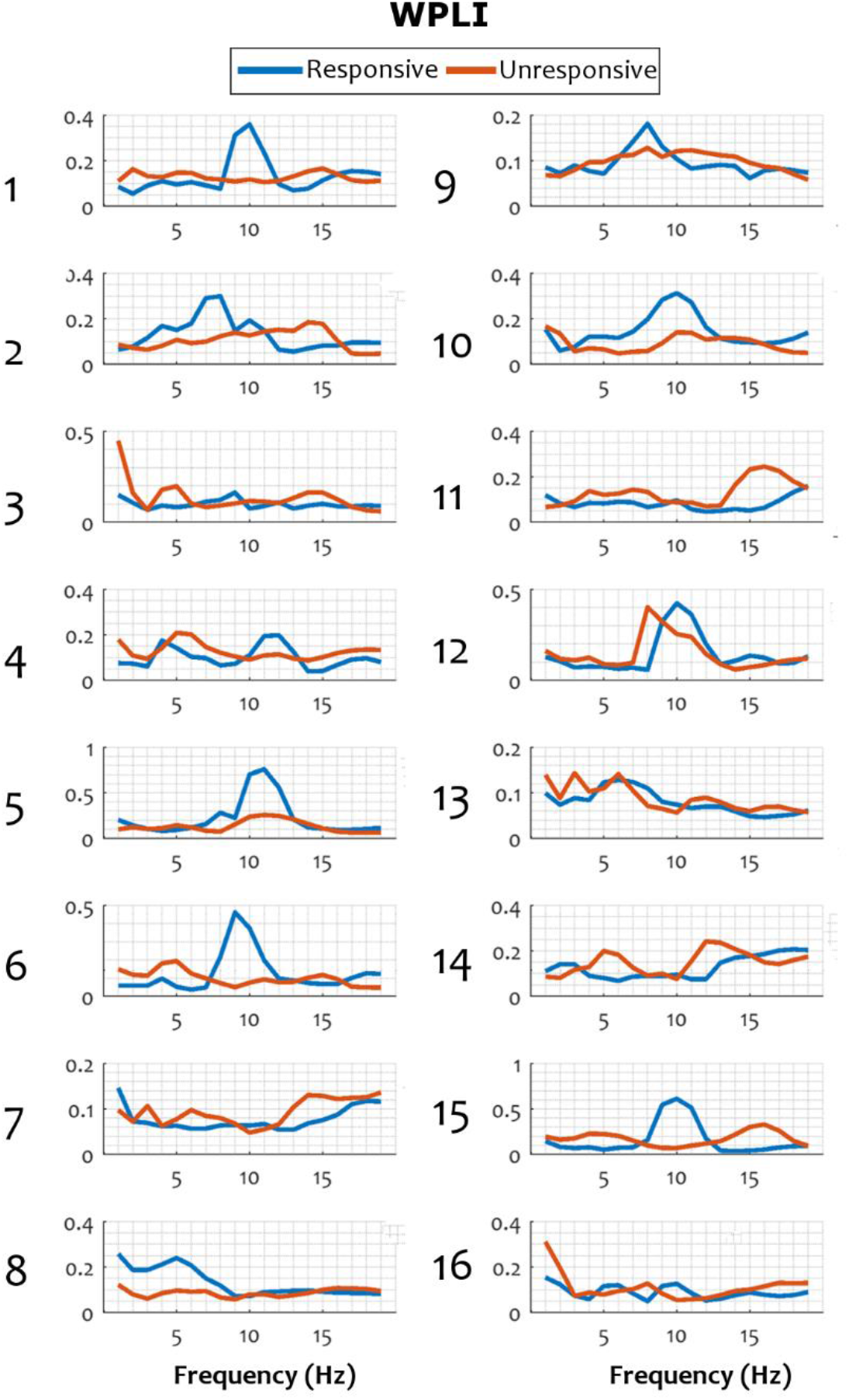
Median WPLI before and after loss of responsiveness due to drowsiness in individual subjects.

**Supplementary Figure 3.**
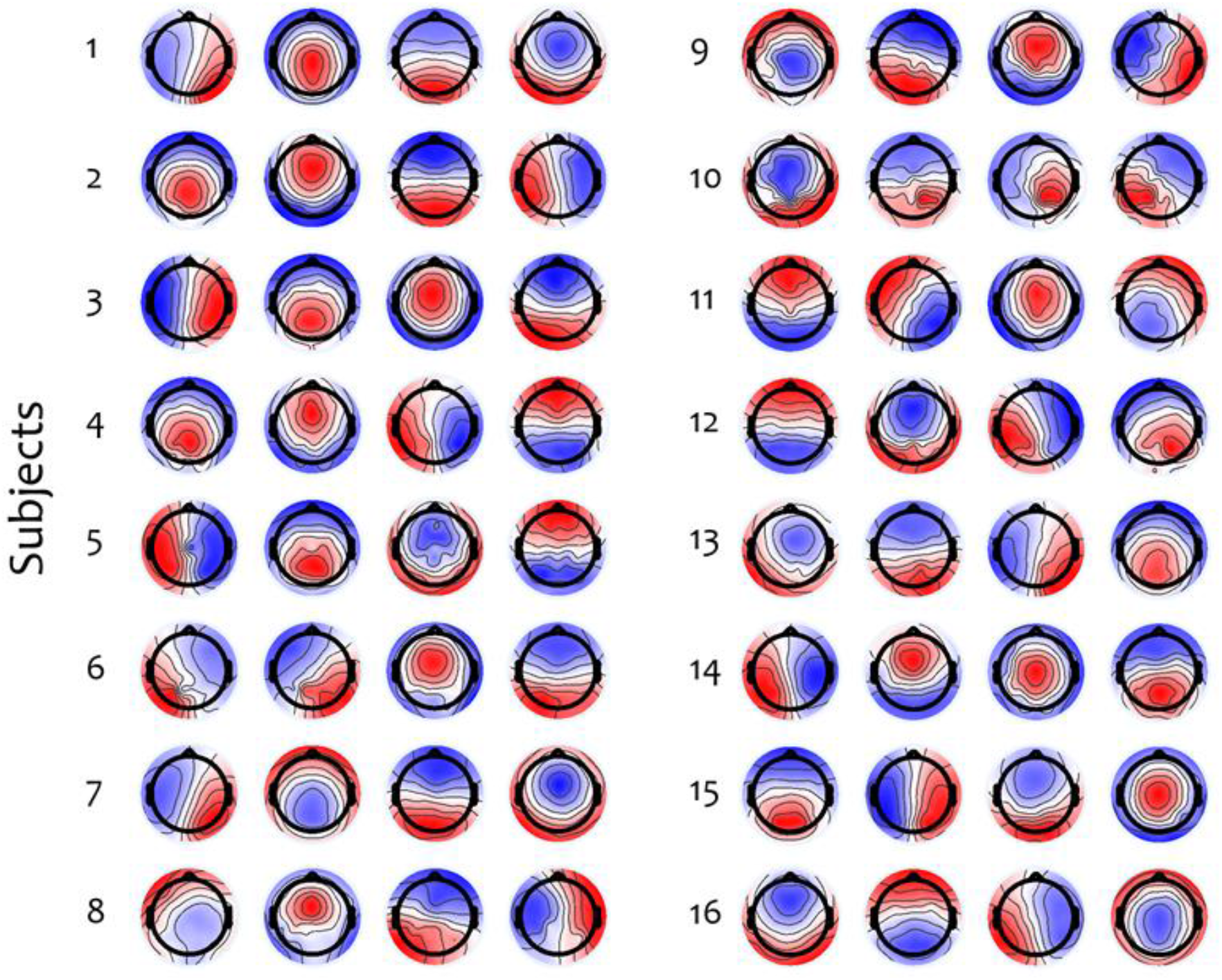
Microstate topographies in each subject, computed over the responsive and unresponsive periods.

